# Multiscale structural control of airflow, diffusion and heat transfer in non-fungus farming termite mounds

**DOI:** 10.64898/2026.04.24.720525

**Authors:** Nengi F Karibi-Botoye, Guy Theraulaz, Bagus Muljadi, Vasily Demyanov, Kamaljit Singh

**Affiliations:** Institute of GeoEnergy Engineering, School of Energy, Geoscience, Infrastructure & Society, Heriot-Watt University, EH14 4AS Edinburgh, United Kingdom; Centre de Recherches sur la Cognition Animale (CRCA), Centre de Biologie Intégrative (CBI), Université de Toulouse, CNRS, UPS, Toulouse, France; Department of Chemical and Environmental Engineering, University of Nottingham, NG7 2RD Nottingham, United Kingdom

**Keywords:** Thermoregulation, ventilation, diffusion, termite mounds, X-ray tomography, numerical simulation, permeability, thermal conductivity, CO_2_ diffusivity, climate control

## Abstract

Termite mounds exhibit highly heterogeneous porous architectures that support efficient ventilation and thermoregulation under variable environmental conditions. Despite considerable interest, particularly in the context of bio-inspired design, the structural properties governing these processes remain only partially understood. In this study, we combine X-ray tomography and numerical simulations to investigate the relationship between structure and climatic properties in non-fungus farming termite mounds built by *Trinervitermes geminatus, Cubitermes* sp., *Apicotermes* sp., and *Thoracotermes* sp. Our results reveal substantial variation in airflow, diffusive transport, and thermal behaviour within mound interiors, driven by differences in internal architecture. At the scale of the whole mound, transport is strongly constrained by the outer walls, which act as the primary resistance to flow and diffusion, thereby largely determining overall permeability and CO_2_ diffusivity. At the microscale, wall microporosity plays a key role in regulating both CO_2_ diffusion and thermal conductivity, with increased microporosity enhancing gas exchange while reducing heat transfer. Within the mound interior, however, structural properties, such as macroporosity, connectivity, and tortuosity, govern transport. Together, these findings demonstrate that transport in termite mounds is inherently multiscale, with microscale wall properties exerting a strong influence on macroscale behaviour. This work advances understanding of the mechanisms underlying passive ventilation and thermoregulation in termite mounds and offers potential guidance for the design of energy efficient, bio-inspired systems.

## 1. Introduction

Termite mounds are complex biogenic structures that have long attracted attention from both scientists and architects due to their remarkable integration of form and function (1). These structures maintain relatively stable thermal, humidity, and ventilation conditions despite strongly fluctuating external environments (2,3). Replicating such passive regulatory mechanisms in human-built systems could improve energy efficiency and reduce CO_2_ emissions by lowering energy demand (4). Achieving this goal, however, requires a detailed understanding of the physical processes governing transport within the mound as well as the structural features that enable and regulate these processes. An interdisciplinary approach combining X-ray tomography and numerical simulations provides deeper insight into the coupling between structure and transport, thereby advancing beyond traditional, predominantly biologically oriented analyses (4).

Termite mounds are constructed from a mixture of soil, saliva, wood fragments, and faecal material (5–8) resulting in mechanically robust structures that can persist for decades to centuries (7,9,10). Termites exhibit high taxonomic diversity, comprising approximately 297 genera and 2951 species (11), and display a wide range of mound architectures that vary in size, geometry, and colony scale (2,12–14). Mounds are broadly classified into fungus farming and non-fungus farming types. Fungus farming species cultivate symbiotic *Termitomyces* fungi and must maintain tightly regulated thermal and humidity conditions to support fungal growth, whereas the fungi contribute to nutrient processing for the colony. In contrast, non-fungus farming termites do not maintain fungal gardens and are therefore subject to different environmental and physiological constraints. This study focuses on non-fungus farming species, specifically *Trinervitermes geminatus, Cubitermes* sp., *Apicotermes* sp., and *Thoracotermes* sp.

Non-fungus farming termite mounds are generally smaller than fungus farming mounds and do not require the maintenance of environmental conditions suitable for fungal growth. At the millimetre scale, these mounds consist of a network of chambers and channels, corresponding to void spaces within the structure, separated by inner and outer walls. At the microscale, the walls are composed of a porous matrix of sand, clay, and mineral components (see figure 3 in Karibi-Botoye *et al*. (4)). Previous studies have characterized these structures using X-ray imaging, graph-based approaches, and in situ measurements (15–19). However, numerical investigations of transport processes in non-fungus farming mounds remain limited. Existing work has largely focused on microscale analyses of the wall structure in *Trinervitermes geminatus* (19), leaving the multiscale coupling between structure and transport at the level of the whole mound largely unexplored.

This study combines X-ray tomography with numerical simulations at multiscale levels to investigate how mound architecture influences diffusion, airflow, and heat transport in non-fungus farming termite. Within the multiscale framework, microscale wall properties are explicitly upscaled and incorporated into millimetre-scale mound geometries enabling the quantification of cross-scale interactions between structural features and transport processes. To our knowledge, this represents the first application of combined millimetre scale and multiscale numerical simulations to these systems. The study addresses two main questions for non-fungus farming termite mounds: (1) what structural features enhance ventilation and thermoregulation at millimetre scale? and (2) how do microscale wall properties influence transport behaviour observed at the millimetre scale?

We first describe the materials and methods used for the study including imaging and numerical simulation approaches. This is followed by the results section, which initially examines transport processes within the inner mound structure and subsequently assesses the influence of outer walls and wall microporosity on airflow and heat transport. The implications of these findings are then discussed in relation to the research questions, followed by a concluding section.

## 2. Materials and Methods

The dataset comprises two mounds of *T. geminatus*, six of *Cubitermes*, two of *Apicotermes*, and three of *Thoracotermes*. These species were selected to enable comparison with previously studied systems. All sampled mounds correspond to epigeal structures, with the exception of *Apicotermes*, which constructs fully subterranean nests. Samples were collected from multiple regions in Africa including Guinea, Senegal, Congo, and Cameroon. The mounds were imaged using X-ray tomography at the millimetre scale, with voxel resolutions ranging from 0.3 to 0.85 mm. The resulting three-dimensional datasets were subsequently used as input geometries for numerical simulations. Detailed descriptions of imaging, data processing, and simulation procedures are provided below.

### 2.1 X-ray tomography and image analysis

#### 2.1.1 Millimetre scale imaging and image processing

The four termite mound species were scanned using a medical X-ray computed tomography (CT) system (Somatom Sensation 16, Siemens, Erlangen, Germany) operated at 120 kV and 150 mA, with voxel sizes ranging from 0.3 to 0.85 mm. Representative three-dimensional reconstructions are shown in Figure 1.

**Figure 1:**
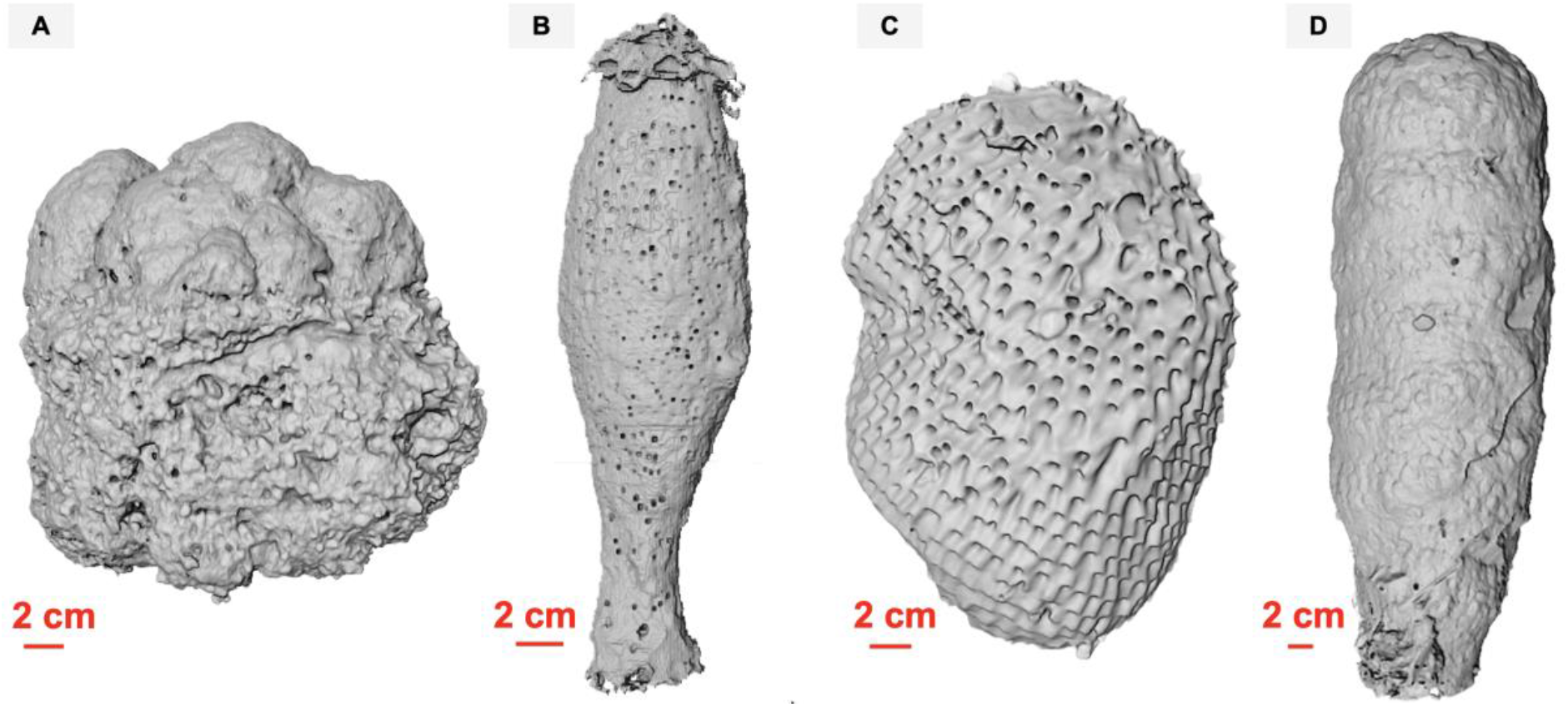
CT reconstructions of non-fungus farming termite mounds. (A) Trinervitermes geminatus mound, (B) Cubitermes sp., (C) Apicotermes sp., (D) Thoracotermes sp. The Apicotermes mound exhibits millimetre-scale openings distributed across the outer wall.

Following acquisition, all datasets were processed to enhance image quality and prepare them for quantitative analysis. Noise reduction was first performed using a non-local means filter (20,21) implemented in Avizo. Image segmentation was then carried out using either watershed methods or the Weka segmentation tool in ImageJ. The images were segmented into solid and air phases with an additional external air phase when present, as described in (4). Subsequent morphological operations, including dilation, erosion, and mask generation, were applied as needed to refine phase boundaries and ensure consistency across datasets. For *Trinervitermes geminatus, Cubitermes*, and *Thoracotermes* mounds only the epigeal portions of the structures were retained for further analysis

Prior to analysis, all images were resampled in Avizo using the Box or Lanczos option to ensure the same voxel size in all three spatial directions as was required by the numerical solver. Differences in porosity between original and resampled images remained below 5%, indicating that the resampling procedure preserved the essential structural features. A representative elementary volume (REV) analysis was performed on the inner regions of mounds (defined here as the internal domains composed of channels and inner walls) prior to numerical simulations. This approach follows the statistical REV (sREV) method in which a cubic window is translated throughout the image while its size is progressively increased until the property of interest converges to a stable value. The properties analysed were the porosity and permeability.

To account for the influence of outer walls on flow and heat transport, artificial outer walls were manually appended to the inner mound sections. This step was necessary because full mound scans included surrounding air, and the geometric complexity of the structures prevented the direct extraction of a computational domain corresponding solely to the mound structure. The outer walls were added along the primary flow direction (x-direction), as illustrated in Figure 2. For the *Apicotermes* mound, millimetre scale holes were incorporated into the outer walls to reproduce their natural architecture. These holes are reported to be small, evenly spaced, and impermeable to termite passage (5). This feature was implemented using hole diameters of less than 3 mm consistent with observational data (Figure 2).

**Figure 2:**
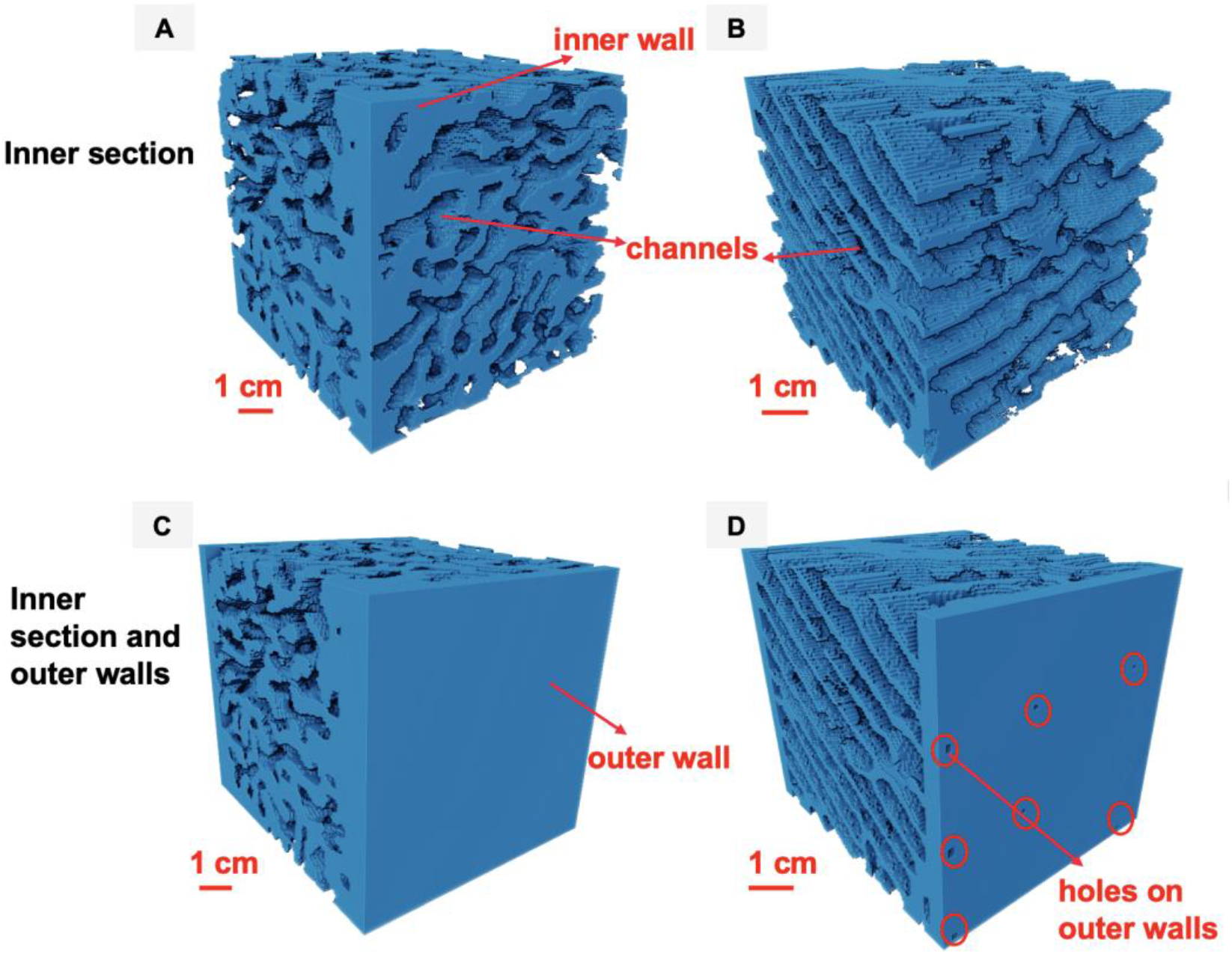
Construction of computational domains by integrating outer walls with inner mound sections for different termite species. (A) Inner section of T. geminatus (B) Inner section of Apicotermes. Inner sections (comprising channels and inner walls) were extended by adding outer wall layers along the principal flow direction (x-direction) to reproduce full mound configurations. Two types of outer wall structures are shown: (C) Non-perforated walls representative of T. geminatus and (D) Perforated walls with millimetre-scale openings representative of Apicotermes. The perforations are regularly distributed and implemented with diameters smaller than 3 mm, consistent with observed structural features.

Another important parameter of the outer walls is their thickness. The ratio of the inner section size and the outer walls thickness was preserved to match that of the full mound, thereby avoiding artefacts in flow and heat transport arising from unrealistic wall thickness. Consequently, outer walls were incorporated into a limited number of subvolumes (two per species), as the procedure was manual and computationally demanding.

#### 2.1.2 Image analysis

Image analysis was performed to quantify key architectural features of the termite mounds, including porosity, height, volume, Euler number, connectivity, and channel thickness. All analyses were conducted using Avizo software. The height and maximum diameter of each sample were measured using the line measurement tool. Porosity was computed as the ratio of air volume to total volume (air and solid phases), based on voxel counts. Tortuosity was determined using the centroid path tortuosity module (22). The integral mean curvature, which quantifies the overall surface curvature of the structure, was computed using the curvature integral module in Avizo. The Euler number, providing a topological measure of connectivity, was computed in 3D as

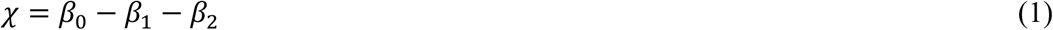

where *β*_*0*_ is the number of connected components, *β*_*1*_ is the number of holes, and *β*_*2*_ is the number of enclosed cavities.

Channel and wall thicknesses were determined by assigning a value of 1 to the phase of interest and the rest of the phases as zero, followed by application of the thickness map module, which computes the diameter of the largest sphere that can be inscribed at each voxel. Connectivity was evaluated using the axis connectivity module, and the fraction of connected structures was quantified as the ratio of connected volume to total volume.

### 2.2 Numerical simulation

Simulations were performed using GeoChemFoam, an open-source solver based on OpenFOAM specifically designed for porous media applications (23). The velocity and pressure fields were obtained by solving the Darcy-Brinkman-Stokes (DBS) equations:

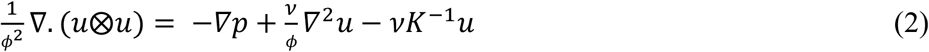

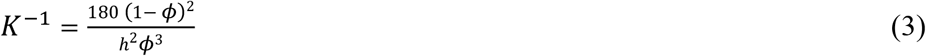

where *u* is the velocity vector, *v* is the kinematic viscosity, *p* is the pressure, *K* is the permeability, *ϕ* is the porosity, and *h* is the mesh resolution.

Air was used as the working fluid, with a kinematic viscosity of 1.602 × 10^−5^ m^2^s^-1^. A volumetric flow rate of 1 × 10^−7^ m^3^s^-1^ was imposed at the inlet of the mound, with zero-gradient boundary conditions at the outlet and no-slip conditions at the solid walls. Under these conditions, the Reynolds numbers remain ≪ 1, corresponding to creeping flow within the Darcy regime and consistent with the low velocities reported in termite mounds (4).

For simulations of pressure, velocity and temperature within the inner mound sections, a structured grid was used. The initial mesh resolution corresponded to approximately half the number of voxels in the image and was subsequently refined once to match the full voxel resolution. This approach reduced computational cost while maintaining accuracy, with deviations in permeability below 5% compared to simulations performed at full resolution.

For multiscale simulations of pressure and velocity, a similar grid strategy was adopted, with one notable exception. For the *Apicotermes* M1-A mound, each voxel was directly mapped to a computational cell of equal size. This higher resolution was required due to the presence of perforated outer walls, as coarser grids resulted in permeability deviations of up to a factor of four. For the *Apicotermes* M4-A mound, however, full-resolution simulations were computationally prohibitive, and the reduced resolution method approach was therefore retained.

These simulations are referred to as multiscale, as effective wall properties (e.g., porosity and permeability) derived from microscale simulations were upscaled and assigned to the walls in the millimetre scale models to represent mound-scale flow. The solid walls of the mounds were assigned a microporosity of 15% and a permeability of 2 × 10^−12^ m^2^, based on microscale simulations reported by Singh *et al*. for *T. geminatus* mounds (19). Cyclic boundary conditions were applied at the inlet and outlet, while zero-gradient conditions were imposed at the walls. The applied pressure drop corresponded to an equivalent flow rate of 1 × 10^−7^m^3^s^-1^.

The permeability across the mounds was determined using Darcy’s equation:

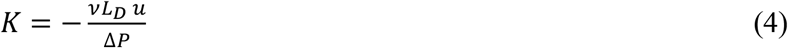

where Δ*P* is the pressure drop between inlet and outlet, *L*_*D*_ is the full length of the domain (23).

Thermal conductivity under steady state is described by

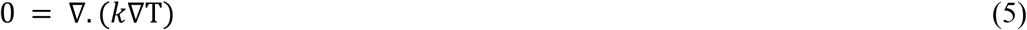

where T is temperature.

The thermal conductivity can be calculated from the harmonic average of thermal conductivity of each phase weighed by its volume fraction (Equation 6) or arithmetic average (Equation 7).

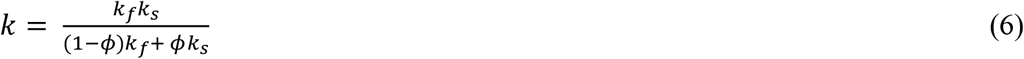

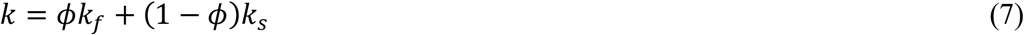

where *k*_*f*_ is thermal conductivity of the fluid and *k*_*s*_ is the thermal conductivity of solid.

Harmonic averaging corresponds to a configuration in which heat transfer occurs sequentially across phases along the direction of heat flow, thereby providing a lower bound for the effective thermal conductivity, whereas arithmetic averaging assumes preferential heat flow through the more conductive phase, and therefore represents an upper bound (24). The solid pore structure of the mound walls is highly heterogeneous and cannot be adequately represented by either idealised configuration alone. Therefore, the effective thermal conductivity was estimated as the mean of the of the harmonic and arithmetic bounds, yielding a value constrained within these limits.

The solid walls were assigned a thermal conductivity of 6.3 W/m·K, a specific heat capacity of 767.64 J/kg·K, and a density of 3003.16 kg/m^3^. These values were derived from the arithmetic weighted average of the mineral composition of the Guinea mound reported by Singh *et al*. (19). Arithmetic averaging was adopted to reflect the high connectivity of the solid phase within the mound structure. For the thermal simulations, the temperature was set to 1 at the inlet and 0 at the outlet, with zero-gradient boundary conditions imposed at the walls. The initial temperature field was uniformly set to 1, and steady-state conditions were assumed.

CO_2_ transport was modelled using Fick’s second law under steady-state conditions:

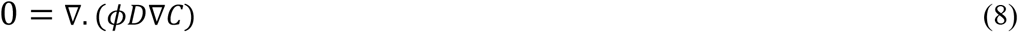

where C is the concentration of the CO_2_ and D is the diffusion coefficient.

For all CO_2_ simulations, except for *Apicotermes* M3_A and M4_A, a structured grid was used in which each voxel was directly mapped to a numerical grid cell of equal size. For *Apicotermes* M3_A and M4_A, a reduced grid resolution corresponding to half the number of voxels was used due to computational constraints. The CO_2_ concentration was set to 1 at the inlet and 0 at the outlet, with zero-gradient boundary conditions at the walls and an initial concentration field set to zero under steady-state conditions. In multiscale simulations, the setField option was used to assign CO_2_ diffusivity values to the outer walls. For all mounds except *Apicotermes*, the outer wall diffusivity was set to 3.1 × 10^−7^ m^2^s^-1^, based on the microscale estimates reported by Singh *et al*. (19). This value was not applied to *Apicotermes*, as it was derived from mounds with microporous but non-perforated walls, which differ structurally from the perforated outer walls observed in this genus.

For the multiple regression analysis of the results, models were evaluated for normality of residuals, homoscedasticity, and multicollinearity. A power-law formulation was adopted for permeability, as it improved predictive performance. For thermal conductivity, surface area was excluded due to its strong correlation with volume, and the Euler number was omitted because it had a negligible effect on *R*^*2*^ and introduced heteroscedasticity.

## 3. Results and discussion

We investigated the mounds of *T. geminatus, Cubitermes, Apicotermes*, and *Thoracotermes*. X-ray imaging of the inner sections reveals marked differences in their internal architectures. *Cubitermes* and *Thoracotermes* mounds are characterized by relatively large, bubble-like chambers, whereas *Apicotermes* and *T. geminatus* are characterized by thinner and more elongated channels (see figure 7A in Karibi-Botoye *et al*. (4)). These architectural differences are associated with variation in chamber size, with larger chambers observed in *Cubitermes* and *Thoracotermes* compared to *Apicotermes* and *T. geminatus* (see figure 7B in Karibi-Botoye *et al*. (4)). Across most mounds, the outer walls are thicker than the inner walls by a factor ranging from 1 to 3.3. In addition, channel dimensions generally exceed inner wall thickness, indicating that void space dominates the internal structure and forms a connected network of pathways for airflow and termite movement within the mound.

### 3.1 Flow and thermal properties in the inner section of the mound

The inner sections of the mound were characterized using a set of structural descriptors, including porosity, volume, tortuosity, channel thickness, inner wall thickness, and Minkowski functionals (volume (M0), surface area (M1), integral mean curvature (M2), and Euler characteristic (M3)). The complete dataset is provided in the supplementary material (Table S1). The inner sections exhibit high channel porosity, ranging from 0.3 to 0.8. *Apicotermes* mounds display the highest porosity, whereas *Cubitermes* exhibits the greatest variability. Channel connectivity exceeds 83% in most cases, with lower values observed in a few *Thoracotermes* mounds that were modified by smaller species, resulting in reduced porosity and connectivity. Velocity fields are spatially heterogeneous within the mounds, reflecting variations in channel architecture (see Figure 3E - 3H).

**Figure 3:**
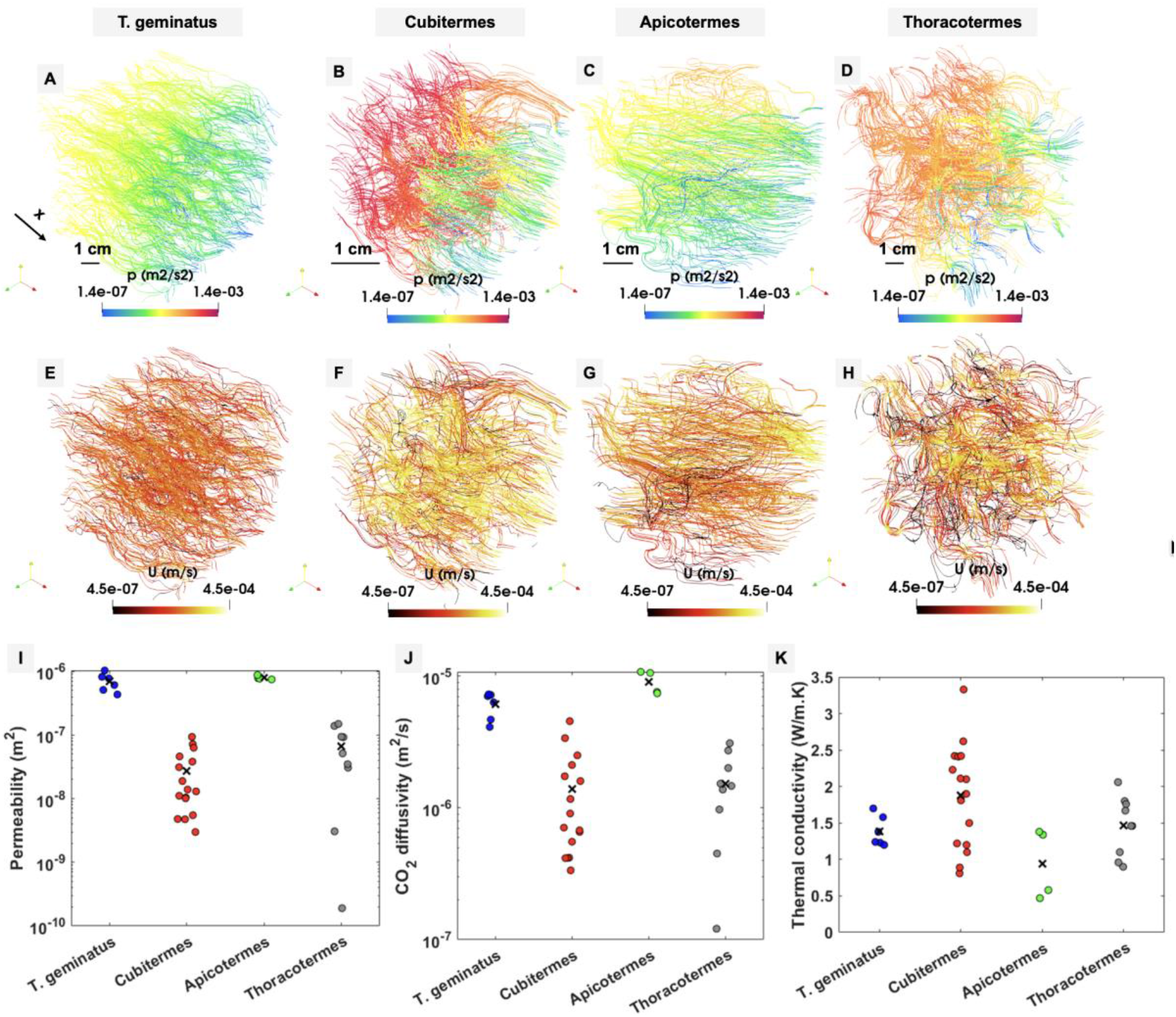
Flow fields and transport properties in the inner sections of non-fungus farming termite mounds. (A to D) Pressure fields and (E to H) Velocity fields for the four mounds species. (I) Permeability, (J) CO_2_ diffusivity, and (K) Thermal conductivity for each species. Blue circles represent Trinervitermes geminatus, red circles Cubitermes, green circles Apicotermes, and grey circles Thoracotermes. Cross symbols indicate the mean value for each species. The arrow shows the direction of flow.

The permeability of the inner sections ranges from 1.9 × 10^−10^ to1.63 × 10^−6^ m^2^ (Figure 3I). The highest permeability values are observed in *T. geminatus* and *Apicotermes*, consistent with their higher porosity, greater connectivity, and lower tortuosity, which together reduce flow resistance. In contrast, *Cubitermes* and *Thoracotermes* exhibit lower permeability. The modified *Thoracotermes* mounds with reduced porosity and connectivity (M4-Th and M5-Th) display permeability values up to two orders of magnitude lower than those of other mounds within the same species.

**Table 1:** Flow, diffusion and thermal properties within the inner section of non-fungus farming mounds.

| Name of species | Subvolume ID | Permeability (m<sup>2</sup>) | Average velocity (m/s) | CO<sub>2</sub> diffusivity (m<sup>2</sup>/s) | Thermal conductivity (W/m.K) |
| --- | --- | --- | --- | --- | --- |
| ***T. Geminatus*** | M1_T | 6.08E-07 | 8.93E-06 | 6.34E-06 | 1.4 |
|  | M2_T | 7.73E-07 | 1.01E-05 | 7.06E-06 | 1.2 |
|  | M3_T | 8.19E-07 | 8.19E-06 | 7.22E-06 | 1.2 |
|  | M4_T | 1.03E-06 | 9.60E-06 | 7.29E-06 | 1.2 |
|  | M5_T | 4.32E-07 | 1.92E-05 | 4.12E-06 | 1.7 |
|  | M6_T | 5.10E-07 | 2.17E-05 | 4.69E-06 | 1.6 |
| ***Cubitermes*** | M1_C | 5.49E-09 | 3.60E-05 | 3.36E-07 | 2.4 |
|  | M2_C | 1.39E-08 | 4.55E-05 | 7.09E-07 | 2.4 |
|  | M3_C | 1.30E-08 | 5.95E-05 | 6.58E-07 | 2.6 |
|  | M4_C | 3.12E-08 | 1.78E-05 | 1.60E-06 | 1.9 |
|  | M5_C | 1.04E-08 | 3.77E-05 | 5.54E-07 | 2.2 |
|  | M6_C | 4.60E-08 | 5.94E-05 | 2.12E-06 | 1.5 |
|  | M7_C | 2.99E-09 | 5.68E-05 | 4.19E-07 | 2.1 |
|  | M8_C | 4.79E-09 | 5.68E-05 | 6.77E-07 | 2.1 |
|  | M9_C | 9.34E-08 | 4.00E-05 | 4.56E-06 | 0.8 |
|  | M10_C | 7.19E-08 | 4.94E-05 | 3.39E-06 | 0.9 |
|  | M11_C | 6.29E-08 | 1.11E-04 | 2.51E-06 | 1.2 |
|  | M12_C | 1.11E-08 | 4.94E-05 | 9.08E-07 | 1.8 |
|  | M13_C | 4.74E-09 | 8.17E-05 | 4.17E-07 | 2.4 |
|  | M14_C | 1.89E-08 | 4.94E-05 | 1.17E-06 | 1.1 |
|  | M15_C | 3.81E-08 | 4.00E-05 | 1.74E-06 | 1.2 |
|  | M16_C | 1.01E-08 | 8.18E-05 | 4.16E-07 | 3.3 |
| ***Apicotermes*** | M1_A | 7.45E-07 | 1.87E-05 | 7.60E-06 | 0.5 |
|  | M2_A | 7.78E-07 | 1.73E-05 | 7.42E-06 | 0.6 |
|  | M3_A | 7.66E-07 | 2.17E-05 | 1.06E-05 | 1.3 |
|  | M4_A | 8.74E-07 | 2.17E-05 | 1.08E-05 | 1.4 |
| ***Thoracotermes*** | M1_Th | 1.39E-07 | 7.55E-05 | 1.53326E-06 | 1.5 |
|  | M2_Th | 3.06E-08 | 8.75E-05 | 1.38E-06 | 1.5 |
|  | M3_Th | 5.13E-08 | 1.92E-05 | 1.47E-06 | 1.8 |
|  | M4_Th | 1.90E-10 | 3.46E-05 | 1.21E-07 | 1.8 |
|  | M5_Th | 3.06E-09 | 3.46E-05 | 4.51E-07 | 1.7 |
|  | M6_Th | 3.47E-08 | 1.76E-05 | 9.74E-07 | 2.1 |
|  | M7_Th | 9.21E-08 | 1.14E-05 | 2.01E-06 | 1.1 |
|  | M8_Th | 1.49E-07 | 1.14E-05 | 3.10E-06 | 0.9 |
|  | M9_Th | 9.36E-08 | 1.03E-05 | 2.73E-06 | 1.0 |

CO_2_ diffusivity ranges from 1.21 × ^10−7^ to 1.08 × 10^−5^ m^2^s^-1^ (Figure 3J) and follows a similar trend to permeability, with the highest values observed in *T. geminatus* and *Apicotermes*. This correspondence likely arises because the same structural features that facilitate fluid flow, such as high porosity, strong connectivity, and low tortuosity also promote CO_2_ transport within the mounds.

Thermal conductivity ranges from 0.5 to 3.3 W/m·K across the mounds and exhibits an inverse relationship with both permeability and CO_2_ diffusivity (Figure 3K). *T. geminatus* and *Apicotermes* display the lowest thermal conductivity values, whereas *Cubitermes* and *Thoracotermes* exhibit higher values. This trend indicates that more porous structures reduce heat transfer through the solid phase, thereby enhancing the insulating properties of the mound.

### 3.2 Influence of structural properties on transport behaviour in the inner section of the mound

To better understand the mechanisms controlling flow, diffusion, and heat transport within the inner mound sections, we examined the influence of key structural properties on these processes.

Porosity shows a generally positive relationship with permeability across most mounds, with higher porosity associated with increased permeability (Figure 4A). However, this relationship is not consistent across all species, indicating that permeability is not governed by porosity alone but also depends on the organisation of the void space. In particular, mounds with similar porosity can display markedly different permeability values due to variations in connectivity and tortuosity.

**Figure 4:**
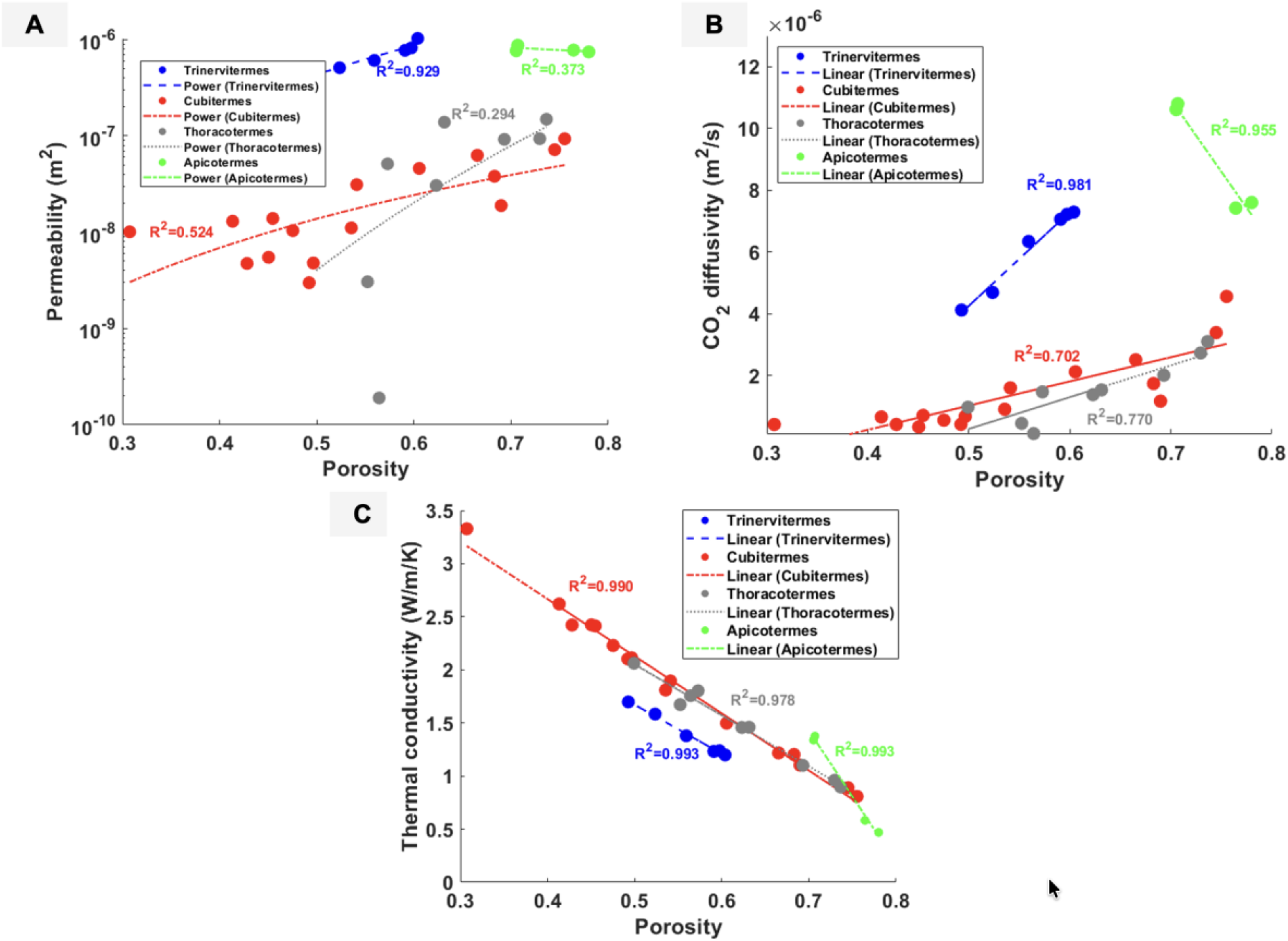
Relationships between porosity and transport properties in different mounds. (A) Porosity–permeability relationship, (B) Porosity–CO_2_ diffusivity relationship, and (C) Porosity–thermal conductivity relationship.

In contrast, CO_2_ diffusivity shows a more consistent dependence on porosity (Figure 4B), with higher porosity leading to increased diffusivity as a result of more continuous and accessible transport pathways. This trend is not observed in *Apicotermes*, which instead shows the opposite trend, potentially due to the different grid setup used for the mounds in this species, which may influence the diffusivity values. Thermal conductivity exhibits the opposite trend, decreasing with increasing porosity (Figure 4C), reflecting the reduced contribution of the solid phase to heat transfer in more porous structures.

While porosity provides a useful first-order descriptor, it does not fully account for the observed variability in transport behaviour within the mounds. To address this limitation, the combined influence of multiple structural properties was assessed using multiple regression analysis. For permeability, the regression model is statistically significant (p < 0.001; F(4,30) = 31.27; R^2^ = 0.81; R^2^adj = 0.78), indicating that permeability is jointly controlled by channel connectivity, porosity, tortuosity, and inner wall thickness (Figure 5A). The adjusted coefficient of determination (R^2^adj) provides a measure of model fit that accounts for the number of predictors, reducing the risk of overfitting. Increased connectivity and porosity enhance permeability, whereas higher tortuosity reduces it by increasing the complexity of flow paths. The positive association between inner wall thickness and permeability likely reflects its correlation with more connected and porous channel networks rather than a direct effect of wall thickness. The equation relating permeability to the independent variables is provided in the supplementary material.

**Figure 5:**
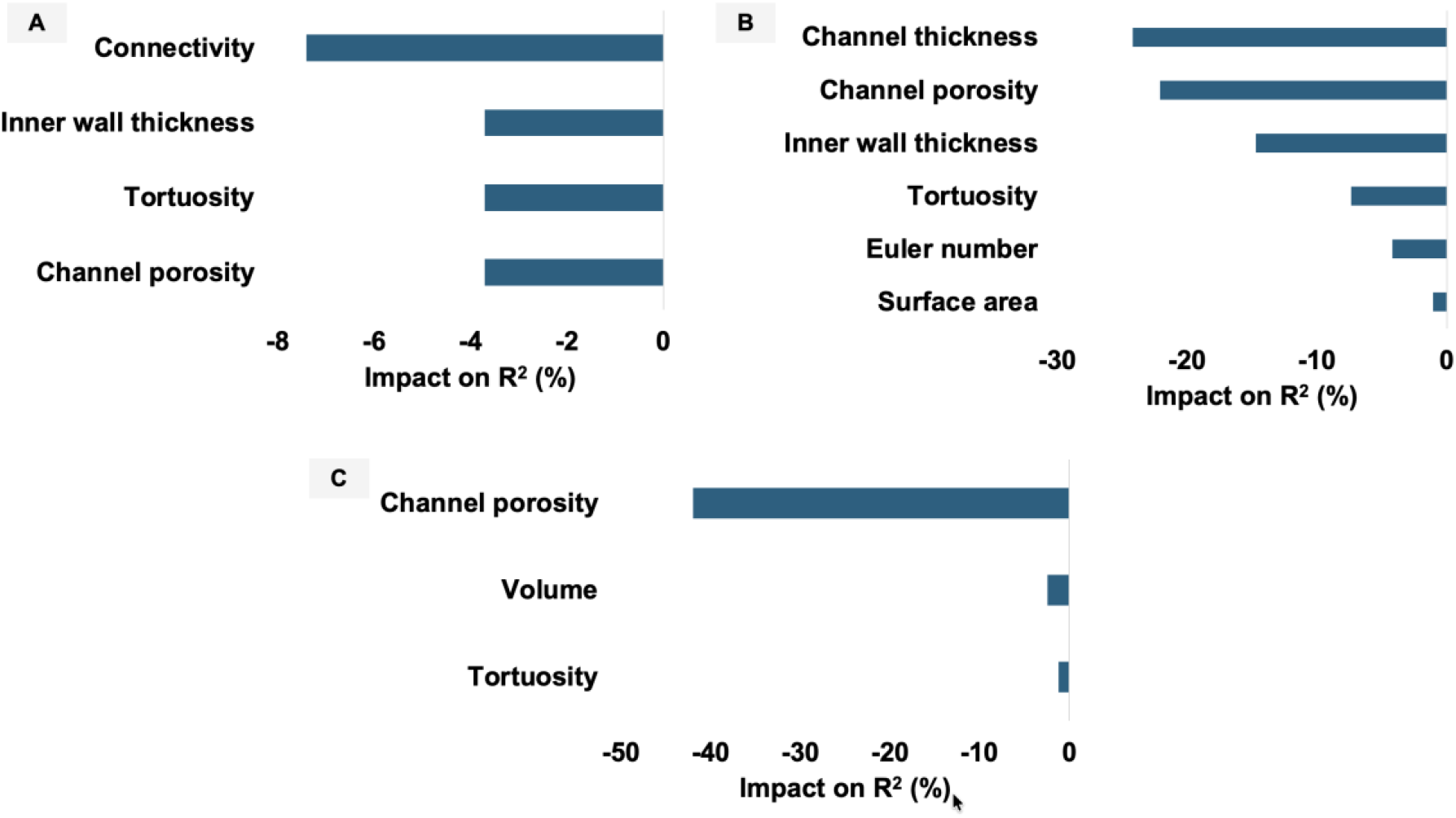
Sensitivity analysis of the coefficient of determination (R^2^) for different mounds. (A) Permeability R^2^ sensitivity, (B) CO_2_ diffusivity R^2^ sensitivity, and (C) Thermal conductivity R^2^ sensitivity.

For CO_2_ diffusivity, the regression model is also highly statistically significant (p < 0.001; F(6,28) = 40.57; R^2^ = 0.90; R^2^adj = 0.87). Channel porosity and channel thickness emerge as the dominant controlling variables, with higher porosity associated with increased diffusivity (Figure 5B). Thinner or more elongated channels are linked to more continuous diffusion pathways, although this relationship is inferred from observed trends rather than directly quantified. A similar pattern is observed for thermal conductivity, for which porosity remains the primary controlling factor, while other structural parameters exert a comparatively weaker influence (Figure 5C). The corresponding regression model is also highly significant (p < 0.001; F(3,31) = 216.68; R^2^ = 0.95; R^2^adj = 0.95). Thermal conductivity decreases with increasing porosity, reflecting enhanced thermal insulation in more porous mound structures.

Overall, these results indicate that transport processes within termite mound interiors are governed by the combined effects of multiple structural properties rather than any single parameter. Porosity controls the available volume for transport, connectivity determines the degree to which these pore spaces are interconnected, and tortuosity controls the efficiency of transport by influencing the complexity of flow and diffusion pathways. Inner wall dimensions further modulate these processes by affecting channel connectivity and local resistance within the structure. Together, these structural characteristics establish the balance between airflow, diffusive transport, and heat transfer within the mound.

### 3.3 Influence of outer walls on diffusion, airflow and heat transport through the mounds

Simulations of the inner sections (channels and inner walls) were extended to include outer walls in order to represent the full mound structure, following the methodology described in section 2. The outer walls were modelled as microporous layers with a porosity of 15%, permeability of 2 × 10^−12^ m^2^ and CO_2_ diffusivity of 3.1 × 10^−7^ m^2^s^-1^, based on Singh *et al*., (19). This approach yields a multiscale representation that couples millimetre scale channels with microporous outer walls layers.

The porosity distributions for *T. geminatus* (with microporous outer walls) and *Apicotermes* mounds (with millimetre-scale wall perforations) are shown in Figure 6A and 6B, respectively. The resulting velocity fields are shown in Figure 6C & 6D. For mounds with microporous outer walls (Figure 6C), the velocity distribution closely resembles that observed in the inner sections without outer walls (see Figure 3E). In contrast, *Apicotermes* mound exhibits more heterogeneous velocity fields due to the presence of millimetre scale holes in the outer walls, which generate localised high velocity regions. The inclusion of outer walls leads to a substantial reduction in overall mound permeability (Figure 6E and Table 2), with decreases of up to five orders of magnitude observed in mounds with microporous walls. Despite significant differences in the permeability of the inner sections, the effective permeability of the full mounds converges to similar values across species, demonstrating that transport at the mound scale is primarily controlled by the outer walls.

**Table 2:** CO_2_ diffusivity, permeability and thermal conductivity in different mounds with outer walls.

| Species | Mound ID | Outer wall thickness (mm) | Length of mound (mm) | Porosity | Permeability (m<sup>2</sup>) | CO<sub>2</sub> diffusivity (m<sup>2</sup>/s) | CO<sub>2</sub> diffusivity (m<sup>2</sup>/s) with micropores of 47% porosity | Thermal conductivity (W/m/K) | Thermal conductivity (W/m/K) with micropores of 47% porosity |
| --- | --- | --- | --- | --- | --- | --- | --- | --- | --- |
| **T. geminatus** | M4-T | 5 | 112 | 0.6 | 2.12E-11 | 5.04E-07 | 1.49E-06 | 0.7 | 0.4 |
|  | M5-T | 14 | 193 | 0.5 | 1.23E-11 | 3.08E-07 | 9.32E-07 | 1.0 | 0.6 |
| **Cubitermes** | M7-C | 2 | 46 | 0.5 | 2.01E-11 | 5.54E-07 | 1.64E-06 | 1.2 | 0.7 |
|  | M15-C | 2 | 55 | 0.7 | 2.12E-11 | 5.65E-07 | 1.67E-06 | 0.7 | 0.4 |
| **Apicotermes** | M1-A | 4 | 81 | 0.7 | 8.74E-11 | 8.57E-06 | 1.34E-05 | 0.3 | 0.2 |
|  | M4-A | 2 | 73 | 0.7 | 3.32E-11 | 9.56E-06 | 1.34E-05 | 0.8 | 0.5 |
| **Thoracotermes** | M7-Th | 3 | 99 | 0.7 | 3.05E-11 | 8.66E-07 | 2.46E-06 | 0.6 | 0.4 |
|  | M1-Th | 3 | 43 | 0.6 | 1.18E-11 | 3.18E-07 | 1.00E-06 | 0.9 | 0.5 |

**Figure 6:**
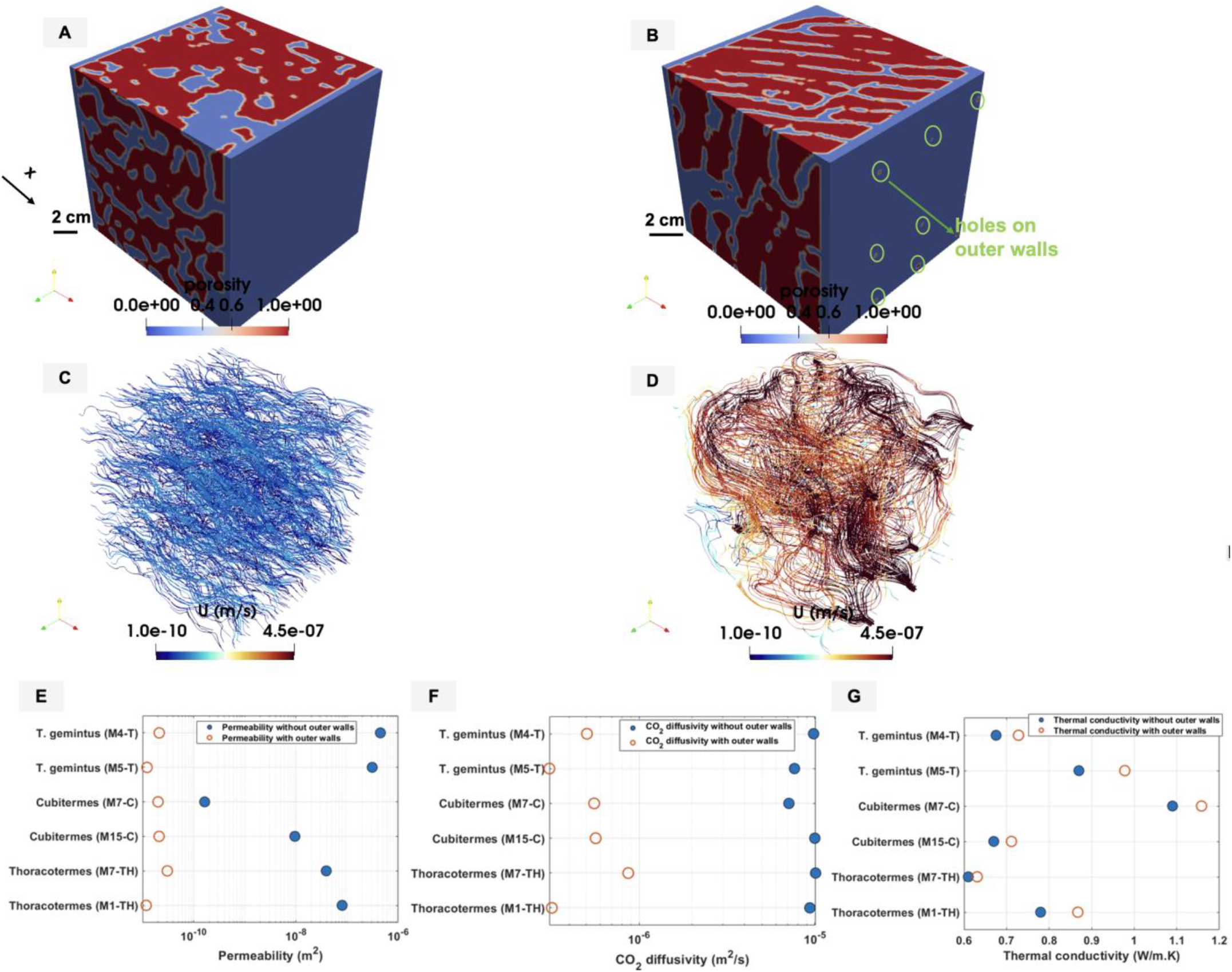
Numerical simulations of multiscale mound structures. (A) Porosity distribution in T. geminatus; (B) Porosity distribution in Apicotermes; (C) Velocity field in T. geminatus; and (D) Velocity field in Apicotermes. (E) Permeability, (F) CO_2_ diffusivity, and (G) Thermal conductivity for different mounds with and without microporous outer walls. When present, both inner and outer walls have a microporosity of 15%.

For mounds with microporous outer walls, the simulated permeability closely agrees with the weighted harmonic average of the inner and outer sections, with values ranging from 0.9 to 1.1 times the predicted values. In contrast, *Apicotermes* mounds with millimetre-scale wall openings deviate from this behaviour, as flow is preferentially channelled through these highly permeable pathways rather than being uniformly distributed across the wall thickness.

Weighted harmonic average permeability (Equation 9) allows the relative contributions of the inner section and outer walls to overall mound permeability to be quantified.

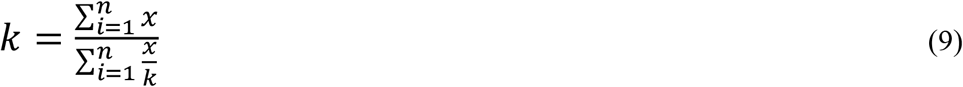

*w*h*ere x is t*h*ickness of t*h*e mound section* and *k*= permeability of that section.

Using the weighted harmonic averaging approach (Equation 9) to calculate overall mound permeability indicates that variations in the permeability of the inner sections have a negligible impact on overall mound permeability of the mound. Increasing the inner section permeability by two orders of magnitude results in only marginal changes. Similarly, reducing the inner section’s permeability by four orders of magnitude, comparable to the values observed in M4-Th and M5-Th from mound 70 constructed by a smaller species, leads to a decrease in overall mound permeability of less than 10%. In contrast, the properties of the outer wall exert a dominant control. Doubling the outer wall permeability from 2 × 10^−12^ to 4 × 10^−12^ m^2^ results in an approximately proportional increase in the overall mound permeability. Similarly, variations in layer thickness also significantly influence mound permeability. Increasing the thickness of the inner section while keeping other parameters constant leads to an increase in overall permeability, with a doubling of inner section thickness resulting in an approximately 1.9-fold increase. A similar effect is observed when the outer wall thickness is reduced. These results demonstrate that mound scale permeability is strongly governed by the properties of the outer walls and the relative thickness of structural layers (Figure 7).

**Figure 7:**
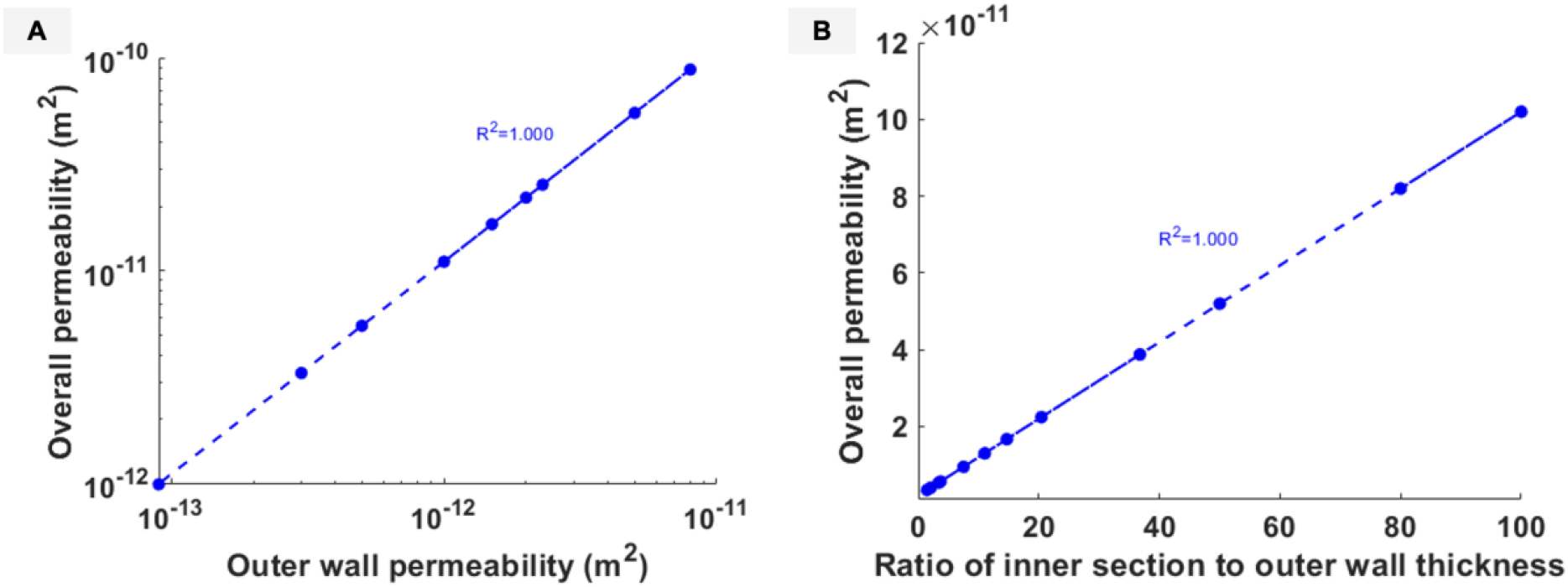
Influence of outer wall properties and structural layer thickness on mound permeability. (A) Effect of outer wall permeability; (B) Effect of the ratio between inner section thickness and outer wall thickness.

A similar trend is observed for CO_2_ diffusivity when outer walls are included. The effective diffusivity is lower than that of the corresponding inner sections, with reductions of up to one order of magnitude (see Figure 6F). CO_2_ diffusivity is strongly controlled by the properties of the outer wall, as the effective diffusivity of the entire mound closely approaches that of the outer walls (3.1 × 10^-7^ m^2^/s). In contrast, *Apicotermes* mounds exhibit higher diffusivity due to the presence of millimetre-scale wall openings in the outer walls, which increase porosity and provide more direct diffusion pathways. This observation is consistent with the environmental conditions in which these mounds are typically found, often in humid soils where ventilation by diffusion is more constrained.

Thermal conductivity of mounds with outer walls ranges from 0.3 to 1.2 W/m·K (see Figure 6G) and exhibits a similar dependence on porosity as observed in the inner sections (see Figure 5C). Although the inclusion of outer walls increases the effective bulk thermal conductivity, the greater wall thickness and associated thermal mass are expected to enhance insulation by slowing heat transfer across the mound.

### 3.4 Influence of increasing microporosity on diffusion, airflow and heat transport in the mounds

In the preceding analysis, a default microporosity of 15% was used for the outer walls of the mounds. However, detailed measurements of wall microporosity are currently available only for *T. geminatus*, and comparable microscale data are lacking for the species. It is therefore important to assess the sensitivity of airflow, diffusion, and heat transfer to variations in wall microporosity. To investigate this effect, the wall microporosity was increased to 47%, corresponding to the upper range of values reported in the literature for fungus farming termite mounds (3,25).

Increasing wall microporosity does not significantly alter the overall permeability of the mound, including in *Apicotermes* mounds that contain millimetre scale openings. This is because the primary factors controlling permeability – namely outer wall permeability, inner section thickness, and outer wall thickness – remain unchanged during the simulations. A similar trend is observed within the inner section of the mound, where permeability remains unchanged under the current assumptions.

In contrast, the CO_2_ diffusivity increases by up to a factor of three as microporosity is raised from 15% to 47% as shown in Figure 8A. The relative enhancement is more pronounced in mounds with lower initial CO_2_ diffusivity, as additional micropores create new pathways for gas transport. This explains why *Apicotermes* mound exhibit a smaller increase (approximately 1.5-fold) as its CO_2_ diffusivity is already high compared to 2.95 – 3.03 -fold increases observed in the other mounds. Increasing microporosity also reduces the thermal conductivity of the mounds, by up to a factor of 1.5, as shown in Figure 8B. This reduction arises from a decrease in the solid phase, which constitutes the primary conducting phase of the mound. Consequently, increasing wall microporosity simultaneously enhances diffusive gas exchange while improving thermal insulation. These effects are particularly advantageous for non-fungus farming termite mounds, where ventilation relies primarily on diffusion (16).

**Figure 8:**
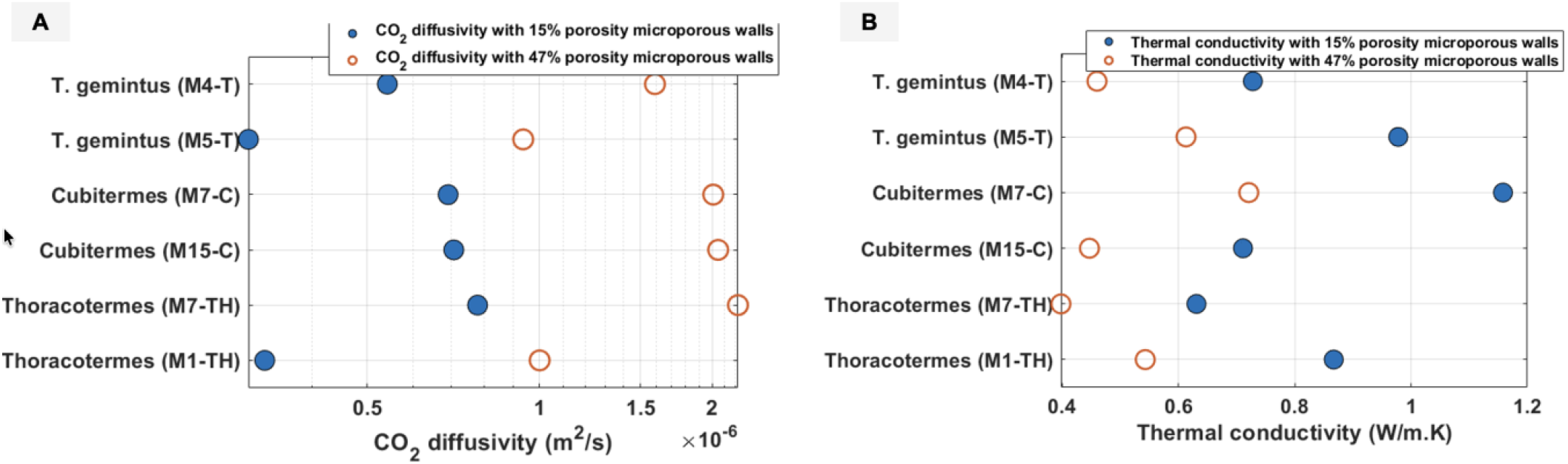
Influence of increasing outer walls microporosity on CO_2_ diffusivity (A) and thermal conductivity (B) in non-fungus farming mounds. The microporosity is increased from 15% to 47% in these simulations.

To address the first research question —namely, which structural features enhance ventilation and thermoregulation at the millimetre (mound) scale – the results show that these processes are governed by structural features operating across multiple scales. At the mound scale, the outer walls act as the primary control layer for ventilation by regulating gas exchange with the external environment. Their effective diffusivity, permeability, and thickness determine the magnitude of gas exchange. Within the mound interior, high channel porosity, strong connectivity, and low tortuosity facilitate internal transport by promoting efficient airflow and heat transfer.

With respect to the second research question, microscale properties exert a dominant influence on millimetre scale transport behaviour by determining the effective properties of the outer walls, which in turn govern overall mound performance. In particular, wall microporosity directly controls CO_2_ diffusivity and thermal conductivity, with higher microporosity enhancing diffusive transport while reducing thermal conductivity. These effects propagate across scales, leading to improved gas exchange and increased thermal insulation at the level of the entire mound.

Despite substantial variations in internal mound structure, the overall permeability is primarily controlled by the outer walls, indicating that microscale wall properties effectively define the boundary conditions for macroscale flow and diffusion. These findings show that transport behaviour in termite mounds is inherently multiscale, with microscale wall properties playing a key role in governing airflow, diffusion, and heat transfer at the mound scale.

## 4. Conclusion

Overall, the properties of the outer walls strongly control the transport of air and CO_2_ in non-fungus farming mounds. Increased outer wall permeability leads directly to higher overall mound permeability, while a reduced outer wall thickness relative to the inner section decreases resistance to transport and enhances flow. A similar relationship is observed for CO_2_ diffusion, where higher outer wall diffusivity results in more efficient gas exchange across the mound. A key question is therefore which specific features most effectively enhance outer wall permeability and CO_2_ diffusivity. For non-fungus farming mounds, where ventilation is predominantly diffusion-driven, particular attention should be given to structural characteristics that maximise CO_2_ diffusivity within the outer walls. Previous work by Singh *et al*. (19) has shown that increasing pore size in the outer walls can enhance CO_2_ diffusivity by up to eight times. Additional structural features may also contribute to enhancing the outer wall permeability and diffusivity and should be further investigated using microscale simulations.

Another important factor influencing CO_2_ diffusivity and thermal conductivity is the amount of micropores within the mound walls. As microporosity increases, CO_2_ diffusivity rises accordingly due to the formation of additional flow pathways. In contrast, thermal conductivity decreases, with values approximately halved for a threefold increase in microporosity, as the fraction of the solid phase – the primary conductive medium – is reduced. These results demonstrate that microscale wall properties exert a strong control on flow, diffusion, and heat transfer observed at the millimetre scale, highlighting the importance of characterizing wall microstructure in greater detail.

This finding does not imply that the inner sections of the mound are negligible. On the contrary, they must maintain sufficiently high permeability and CO_2_ diffusivity to ensure efficient internal transport. Such properties can be achieved through increased channel connectivity and porosity combined with reduced tortuosity within the internal network. At the same time, maintaining relatively low thermal conductivity is advantageous for insulation, which is also strongly influenced by porosity.

Taken together, these results demonstrate that ventilation and heat transfer in non-fungus farming mounds are not governed by a single structural parameter, but rather emerge from the combined effects of outer wall properties, wall microporosity, and internal architecture. A natural extension of this work is to compare flow and thermal properties in fungus farming termite mounds across both micro and mound scale levels, in order to identify key similarities and differences with non-fungus farming systems.

This study advances the understanding of ventilation and thermoregulation in termite mounds by explicitly linking structural features across multiple scales. More broadly, the findings highlight that transport processes in biological porous systems are inherently multiscale, and that elucidating the relationship between microscale structure and macroscale performance is essential not only for termite mounds but also for the design of natural and engineered porous materials.

## Acknowledgment

N.F.K. acknowledges Julien Maes and Hannah Menke for their help with the numerical simulations. N.F.K. and K.S. acknowledge funding from the James Watt Scholarship from Heriot-Watt University. N.F.K. acknowledges funding from Mary Burton funds from Heriot-Watt University. G.T. acknowledges funding from Agence Nationale de la Recherche (ANR-06-BYOS-0008) for X-ray medical scanning. G.T. also gratefully acknowledges the Indian Institute of Science to serve as Infosys visiting professor at the Centre for Ecological Sciences in Bengaluru, India.

## Notes

### Competing Interest Statement

The authors have declared no competing interest.

