## Supplementary material (Table S1). for "Multiscale structural control of airflow, diffusion and heat transfer in non-fungus farming termite mounds"

Table S.1: Structural properties of the different mound subvolumes obtained from image analysis. The definitions and methods used to compute properties such as Euler number, channel connectivity, and tortuosity are described in Section 3. Here, connectivity refers specifically to channel connectivity at the millimetre scale

| Name of species | Subvolume ID | Channel Porosity | Length in x direction (m) | Volume ($m^3$) | Average channel thickness (m) | Average inner wall thickness (m) | Channel connectivity | Euler number | Surface area ($m^2$) | Integral mean curvature | Tortuosity |
|---|---|---|---|---|---|---|---|---|---|---|---|
| ***T. Geminatus*** | M1-T | 0.6 | 0.11 | 1.1E-03 | 5.2E-03 | 6.9E-03 | 1.00 | -1345 | 0.31 | 1.15E+04 | 1.65 |
| | M2-T | 0.6 | 0.13 | 1.3E-03 | 5.5E-03 | 5.3E-03 | 1.00 | -1358 | 0.33 | 1.01E+04 | 1.51 |
| | M3-T | 0.6 | 0.11 | 1.4E-03 | 5.7E-03 | 4.5E-03 | 1.00 | -1424 | 0.34 | 9.62E+03 | 1.53 |
| | M4-T | 0.6 | 0.1 | 1.1E-03 | 5.9E-03 | 3.9E-03 | 1.00 | -1201 | 0.28 | 5.99E+03 | 1.52 |
| | M5-T | 0.5 | 0.16 | 8.6E-04 | 6.8E-03 | 9.2E-03 | 1.00 | -651 | 0.18 | 5.14E+04 | 1.74 |
| | M6-T | 0.5 | 0.12 | 5.4E-04 | 6.9E-03 | 7.2E-03 | 1.00 | -457 | 0.12 | 3.71E+04 | 1.63 |
| ***Cubitermes*** | M1-C | 0.5 | 0.05 | 1.5E-04 | 6.5E-03 | 6.2E-03 | 0.91 | 8 | 0.04 | 3.46E+04 | 1.67 |
| | M2-C | 0.5 | 0.05 | 1.0E-04 | 6.6E-03 | 5.1E-03 | 0.84 | 39 | 0.03 | 2.63E+04 | 1.8 |
| | M3-C | 0.4 | 0.04 | 6.9E-05 | 5.0E-03 | 5.5E-03 | 0.97 | 19 | 0.02 | 2.07E+04 | 1.88 |
| | M4-C | 0.5 | 0.04 | 2.5E-04 | 5.8E-03 | 3.7E-03 | 0.97 | -145 | 0.09 | 7.62E+04 | 1.92 |
| | M5-C | 0.5 | 0.05 | 1.4E-04 | 5.7E-03 | 4.0E-03 | 0.90 | -32 | 0.05 | 4.60E+04 | 1.84 |
| | M6-C | 0.6 | 0.04 | 6.9E-05 | 5.9E-03 | 2.8E-03 | 1.00 | -58 | 0.03 | 2.45E+04 | 1.67 |
| | M7-C | 0.5 | 0.04 | 7.4E-05 | 4.1E-03 | 2.3E-03 | 0.98 | 1 | 0.03 | 1.15E+05 | 1.47 |
| | M8-C | 0.5 | 0.04 | 7.4E-05 | 4.0E-03 | 2.4E-03 | 0.99 | -17 | 0.03 | 1.18E+05 | 1.52 |
| | M9-C | 0.8 | 0.05 | 1.3E-04 | 7.0E-03 | 1.9E-03 | 1.00 | -797 | 0.05 | 6.26E+04 | 1.32 |

| Name of species | Subvolume ID | Channel Porosity | Length in x direction (m) | Volume ($m^3$) | Average channel thickness (m) | Average inner wall thickness (m) | Channel connectivity | Euler number | Surface area ($m^2$) | Integral mean curvature | Tortuosity |
|---|---|---|---|---|---|---|---|---|---|---|---|
| | M10-C | 0.7 | 0.05 | 9.1E-05 | 7.0E-03 | 1.3E-03 | 0.98 | -240 | 0.04 | 5.44E+04 | 1.24 |
| | M11-C | 0.7 | 0.03 | 2.7E-05 | 6.4E-03 | 2.4E-03 | 0.99 | 4 | 0.01 | 2.14E+04 | 1.77 |
| | M12-C | 0.5 | 0.05 | 9.1E-05 | 4.4E-03 | 2.3E-03 | 0.95 | -90 | 0.04 | 7.21E+04 | 1.66 |
| | M13-C | 0.4 | 0.04 | 4.3E-05 | 4.0E-03 | 2.9E-03 | 0.93 | 10 | 0.02 | 4.09E+04 | 1.41 |
| | M14-C | 0.7 | 0.05 | 9.1E-05 | 7.6E-03 | 1.9E-03 | 0.97 | -24 | 0.04 | 1.35E+04 | 1.34 |
| | M15-C | 0.7 | 0.05 | 1.3E-04 | 7.7E-03 | 2.1E-03 | 1.00 | -21 | 0.05 | 1.62E+04 | 1.66 |
| | M16-C | 0.3 | 0.04 | 4.3E-05 | 6.5E-03 | 6.8E-03 | 0.67 | 34 | 0.01 | 5.23E+03 | 1.74 |
| ***Apicotermes*** | M1-A | 0.8 | 0.07 | 3.9E-04 | 6.2E-03 | 3.1E-03 | 1.00 | -1112 | 0.12 | 1.24E+04 | 1.09 |
| | M2-A | 0.8 | 0.08 | 4.4E-04 | 5.8E-03 | 2.0E-03 | 1.00 | -1364 | 0.14 | 5.91E+03 | 1.11 |
| | M3-A | 0.7 | 0.07 | 3.1E-04 | 4.4E-03 | 1.6E-03 | 1.00 | 710 | 0.13 | 3.50E+05 | 1.05 |
| | M4-A | 0.7 | 0.07 | 3.1E-04 | 4.4E-03 | 1.6E-03 | 1.00 | 529 | 0.13 | 3.66E+05 | 1.04 |
| ***Thoracotermes*** | M1-Th | 0.6 | 0.04 | 4.8E-05 | 8.7E-03 | 2.6E-03 | 0.96 | 18 | 0.02 | 2.28E+04 | 1.43 |
| | M2-Th | 0.6 | 0.03 | 3.9E-05 | 8.4E-03 | 2.9E-03 | 0.94 | 17 | 0.01 | 1.81E+04 | 1.33 |
| | M3-Th | 0.6 | 0.03 | 2.3E-05 | 7.5E-03 | 3.4E-03 | 0.99 | 18 | 0.05 | 1.63E+04 | 1.5 |
| | M4-Th | 0.6 | 0.05 | 1.6E-04 | 5.6E-03 | 2.6E-03 | 0.23 | 117 | 0.06 | 2.35E+05 | 2.11 |
| | M5-Th | 0.6 | 0.05 | 1.6E-04 | 5.2E-03 | 2.4E-03 | 0.45 | 27 | 0.07 | 2.50E+05 | 2 |
| | M6-Th | 0.5 | 0.08 | 4.3E-04 | 8.1E-03 | 4.1E-03 | 0.96 | -69 | 0.11 | 1.53E+05 | 1.41 |
| | M7-Th | 0.7 | 0.09 | 8.2E-04 | 9.8E-03 | 2.4E-03 | 0.99 | -775 | 0.23 | 2.22E+05 | 1.22 |
| | M8-Th | 0.7 | 0.09 | 8.2E-04 | 9.9E-03 | 2.1E-03 | 0.99 | -1287 | 0.24 | 1.97E+05 | 1.23 |
| | M9-Th | 0.7 | 0.1 | 9.7E-04 | 9.8E-03 | 2.0E-03 | 1.00 | -1121 | 0.29 | 2.54E+05 | 1.24 |

The relationships between structural properties and transport behaviour in the inner sections were quantified using multiple regression analysis, leading to the following empirical correlations:

$$Permeability = 4804.8 \times Porosity^{8} \times Inner\ wall\ thickness^{3.2} \times Connectivity^{2.2} \times Tortuosity^{-4.3} \qquad \text{(S.1)}$$

$$CO_2\ diffusivity = 2.3 \times 10^{-5} \times Channel\ Porosity - 4 \times 10^{-6} \times Tortuosity - 1.1 \times 10^{-3} \times Channel\ thickness + 1.0 \times 10^{-3} \times Inner\ wall\ thickness + 1.6 \times 10^{-9} \times Euler\ number + 1.5 \times 10^{-5} \times Surface\ area - 2.2 \times 10^{-6} \qquad \text{(S.2)}$$

$$Thermal\ conductivity = 5.2 - 0.3 \times Tortousity - 5.3 \times Channel\ porosity - 329.3 \times Volume \qquad \text{(S3)}$$
